# A high throughput deep amplicon sequencing method to show the emergence and spread of *Calicophoron daubneyi* in United Kingdom cattle herds

**DOI:** 10.1101/521229

**Authors:** Neil D. Sargison, Kashif Shahzad, Stella Mazeri, Umer Chaudhry

## Abstract

The prevalence of *C. daubneyi* infection in the United Kingdom has increased, but despite the potential for rumen flukes to cause production loss in ruminant livestock, understanding of their emergence and spread is poor. Here we describe the development of a method to explore the multiplicity of *C. daubneyi* infection and patterns of the parasite’s emergence and spread, based on Illumina MiSeq deep sequencing of meta barcoded amplicons of a fragment of the mt-COX-1 locus. Our results show high levels of genetic diversity per infection and between populations of 10 to 47 of adult *C. daubneyi*, each from a total of 32 finished prime cattle consigned to slaughter from northern United Kingdom; with 18 unique mt-COX-1 haplotypes. This has implications for the adaptability of environmental and intermediate host stages of the parasite to changing climatic and animal management conditions, or of parasitic stages to exposure to anthelmintic drugs; potentially allowing for greater pathogenicity, or the development of anthelmintic resistance, respectively. Our results illustrate the impact of high levels of animal movements in the United Kingdom, whereby multiple common mt-COX-1 haplotypes were identified in 26 populations in the absence of geographical clustering of clades.

## 1. Introduction

Digenean flukes belonging to the family Paramphistomidae are common parasites of livestock that are kept in a wide geographical range of environments supporting the intermediate mud, or water snail hosts (Huson et al., 2017). They are regarded as causes of severe disease and production loss in tropical and subtropical regions (Rolfe et al., 1991; Rangel-Ruiz et al., 2003). Historically, paramphistomes were considered to be uncommon in temperate regions (Wilmott, 1950; Deiana and Arru, 1963), but over the past two decades rumen flukes, now known to be *Calicophoron daubneyi* (Gordon et al., 2013), have become commonplace in wetter parts of Europe, notably in Spain and Portugal (Diaz et al., 2007; Arias et al., 2011; González-Warleta et al., 2013; Ferreras et al., 2014), France (Abrous et al., 2000; Mage et al., 2002; Rieu et al., 2007), Belgium (Malrait et al., 2015), Ireland (Murphy et al., 2008; Zintl et al., 2014; Toolan et al., 2015; Martinez-Ibeas et al., 2016) and the United Kingdom (Foster et al., 2008; Millar et al., 2012; Jones et al., 2017). Prior to the mid 2000s, there were neither anecdotal reports of adult rumen flukes in cattle or sheep forestomachs, nor evidence of paramphistome eggs in faecal samples in the United Kingdom and Republic of Ireland; suggesting the new introduction and subsequent spread of the parasites. Pathogenesis and production loss due to *C. daubneyi* in Europe is mostly associated with the activity of immature flukes in the intestine (Foster et al., 2008; Mason et al., 2012), while there is little compelling evidence for adult flukes in the forestomach causing production loss (Sargison et al., 2016); putatively associated with a switch from feeding on host tissue in the intestine to microbial contents of the forestomach (reviewed by Huson et al., 2017).

A number of factors may have influenced the emergence and spread of *C. daubneyi* infection in Europe, such as: the abundance of suitable *Galba truncatula* intermediate hosts (Abrous et al., 1999; Jones et al., 2017); climatic change favouring the completion of the parasite’s lifecycle (Skuce et al., 2013); the intensity of infection; and movement of free-living or parasitic stages between regions (Blouin et al., 1995). However, despite the pervasiveness of *C. daubneyi*, these epidemiological factors are poorly understood.

Abattoir-based studies have been used to explore the phylogenetics of *C. daubneyi* in cattle in the Republic of Ireland and in Northern Ireland, showing high levels of genetic diversity in the two countries and no evidence of geographical clustering of mitochondrial cytochrome c oxidase subunit I (mt-COX-1) DNA haplotypes (Zintl et al., 2014). The Median Joining Network had a linear rather than a star-like appearance, with the most commonly identified haplotype resembling the only contemporary NCBI reference published sequence (accession number: JQ815200), originating from Spain (Martínez-Ibeas et al., 2013). This implied either recent expansion of a small dispersed established population, or more probably multiple introductions from a larger mainland European parent population. These studies were based on generating individual two direction Sanger sequence data from 885 bp mt-COX-1 DNA fragments (Martínez-Ibeas et al., 2013) derived from 97 flukes (25 each from two farms and 20 from other cattle from the Republic of Ireland and 27 from other cattle presumed from Northern Ireland). Similar approaches, using high throughput technologies to allow the study of a larger and more diverse fluke population could potentially show if *C. daubneyi* infection emerged in United Kingdom cattle at a single time in a single cattle population, or repeatedly in different cattle populations, before spreading as a result of animal movement.

In this paper we describe a study using *C. daubneyi* adult parasites collected from prime cattle slaughtered in a large red meat abattoir, with the aims of: i) confirming the species identity of recovered rumen flukes; ii) identifying the presence of multiple genotypes per infection (multiplicity of infection) and iii) demonstrating the emergence and spread of *C. daubneyi* haplotypes. The species identity of rumen flukes was confirmed by deep amplicon sequencing of a 282 bp fragment of second internal transcribed spacer (ITS-2) rDNA. Haplotype diversity in 32 *C. daubneyi* populations, each derived from different single infected cattle consigned from different locations in the northern United Kingdom was shown by deep amplicon sequencing of a 333 bp fragment of the mt-COX-1 locus. Split and network trees of the mt-COX-1 haplotypes were examined to show patterns of emergence and spread of *C. daubneyi*, providing proof of concept for a novel approach to the study of parasite epidemiology.

## 2. Materials and methods

### 2.1. Study resources

Populations of adult *C. daubneyi* were collected from a total of 32 finished prime cattle slaughtered during 2014 in a large red meat abattoir in central Scotland during the winter (on 5 sampling days between 13^th^ January and 3^rd^ March 2014) and summer/autumn (on 5 sampling days between 25^th^ August and 6^th^ October 2014). The sample collection strategy was designed both for logistical reasons, and to reduce the risk of disproportionate collection of multiple *C. daubneyi* populations from the same holdings, thereby allowing a representative distribution of cattle origins of consignment. All cattle slaughtered in United Kingdom abattoirs are uniquely identified and traceable under the auspices of the British Cattle Movement Services (https://www.gov.uk/cattle-tracing-online), showing each animal’s origin prior to slaughter. Briefly, the identification of every tenth animal slaughtered on each sampling day was checked and recorded at the point of slaughter and the corresponding forestomachs were tagged to allow samples to be collected from the correct animals and matched to BCMS data. Forestomachs were incised along the greater curvature of the rumen and everted to remove their contents, as a standard part of the abattoir’s tripe preparation process. The total numbers of adult *C. daubneyi* were enumerated and between 10 and 50 rumen flukes from each parasitised animal, depending on the numbers of flukes present (Supplementary Table 1), were fixed in 70% ethanol and archived.

**Table 1:** Summary of genetic diversity of mt-COX-1 haplotypes identified from thirty-two opulations of *C. daubneyi* in the United Kingdom.

| Populations ID | Total number of Illumina MiSeq Reads | Total no of haplotype generated from each population | Haplotype diversity ($H_d$) | Segregating sites (S) | Nucleotide diversity ($\Pi$) | Mutation parameter based on S ($\Theta_s$) | Mean number of pairwise difference (k) |
| --- | --- | --- | --- | --- | --- | --- | --- |
| Pop1 | 1128 | 2 | 0.857 | 26 | 0.029 | 0.024 | 9.77 |
| Pop2 | 1081 | 3 | 0.800 | 5 | 0.006 | 0.004 | 2.13 |
| Pop3 | 2166 | 5 | 0.960 | 15 | 0.011 | 0.012 | 3.62 |
| Pop4 | 1697 | 4 | 0.824 | 8 | 0.007 | 0.005 | 2.76 |
| Pop5 | 1551 | 4 | 0.771 | 9 | 0.006 | 0.004 | 3.17 |
| Pop6 | 6600 | 3 | 0.913 | 13 | 0.008 | 0.010 | 2.79 |
| Pop7 | 3840 | 6 | 0.948 | 26 | 0.016 | 0.012 | 4.18 |
| Pop8 | 2232 | 1 |  |  | N/A |  |  |
| Pop9 | 3668 | 6 | 0.965 | 21 | 0.012 | 0.022 | 4.08 |
| Pop10 | 1675 | 3 | 0.824 | 10 | 0.008 | 0.003 | 4.31 |
| Pop11 | 1419 | 5 | 0.952 | 16 | 0.011 | 0.012 | 3.79 |
| Pop12 | 1381 | 3 | 0.824 | 8 | 0.008 | 0.007 | 2.94 |
| Pop13 | 3896 | 6 | 0.977 | 34 | 0.021 | 0.021 | 4.22 |
| Pop14 | 1819 | 2 | 0.812 | 7 | 0.006 | 0.005 | 2.26 |
| Pop15 | 1783 | 3 | 0.935 | 43 | 0.016 | 0.013 | 5.29 |
| Pop16 | 986 | 1 |  |  | N/A |  |  |
| Pop17 | 1824 | 4 | 0.957 | 16 | 0.012 | 0.013 | 4.01 |
| Pop18 | 1039 | 3 | 0.913 | 9 | 0.006 | 0.007 | 2.21 |
| Pop19 | 1190 | 1 |  |  | N/A |  |  |
| Pop20 | 1955 | 1 |  |  | N/A |  |  |
| Pop21 | 1085 | 1 |  |  | N/A |  |  |
| Pop22 | 1210 | 4 | 0.815 | 9 | 0.011 | 0.007 | 3.62 |
| Pop23 | 1162 | 4 | 0.842 | 7 | 0.008 | 0.006 | 2.86 |
| Pop24 | 1213 | 6 | 0.848 | 10 | 0.009 | 0.007 | 3.00 |
| Pop25 | 1903 | 2 | 0.395 | 4 | 0.004 | 0.003 | 1.57 |
| Pop26 | 1397 | 2 | 0.589 | 6 | 0.007 | 0.005 | 2.52 |
| Pop27 | 1176 | 4 | 0.765 | 6 | 0.006 | 0.003 | 2.52 |
| Pop28 | 3396 | 3 | 0.812 | 8 | 0.008 | 0.006 | 2.66 |
| Pop29 | 1911 | 2 | 0.771 | 36 | 0.007 | 0.004 | 2.44 |
| Pop30 | 4477 | 4 | 0.754 | 7 | 0.006 | 0.004 | 2.05 |
| Pop31 | 1090 | 2 | 0.692 | 5 | 0.007 | 0.003 | 2.30 |
| Pop32 | 1993 | 5 | 0.912 | 12 | 0.012 | 0.010 | 3.94 |
| **Total** | **64943** | **18** | **0.949** | **76** | **0.011** | **0.038** | **3.597** |

### 2.2. Genomic DNA extraction

In order to avoid contamination with any progeny DNA, a ∼2 mg section of each parasite’s head was taken and rinsed twice for 5 min in a petri dish in ddH_2_O. The tissue sections were then lysed in lysis buffer and Protinease K (10mg/ml, New England BioLabs). The lysis buffer contained 50 mM KCL, 10 mM Tris (pH 8.3), 2.5 mM MgCl_2_, 0.045% Nonidet p-40, 0.45% Tween-20, 0.01% gelatin and ddH_2_O in 50ml volumes (Chaudhry et al., 2015a). Samples were lysed in 50 μl for 98 minutes at 60°C followed by 15 minutes at 94°C then stored at -80°C until PCR.

### 2.3. Confirmation of fluke DNA

1 μl of each lysate containing fluke DNA was used to make population pools representing between 10 and 47 parasites from each of the sampled cattle. A final dilution of 1:20 of pooled lysate: ddH_2_O was used as PCR template.

### 2.4. Deep amplicon sequencing of r-ITS-2 and mt-COX-1 DNA

Deep amplicon sequencing was performed from a 282 bp fragment of ITS-2 rDNA using previously published universal trematode primer sets (Adlard et al., 1993; Chaudhry et al., 2015a) and from a 333 bp fragment of mt-COX-1 using newly developed primers (Supplementary Table 2A) amplifying a locus within the 885 bp fragment used by Zintl et al. (2014). Adapters were added to allow the successive annealing of the primers and N is the number of random nucleotides included between each specific primer to increase the variety of generated amplicons. Four forward and four reverse primers were mixed in equal proportion in the first round PCR reaction made under the following conditions: 5X KAPA HiFi Fidelity buffer, 10 mM dNTPs, 10 μM forward and reverse adapter primer, 0.5 U KAPA HiFi Fidelity Polymerase (KAPA Biosystems, USA), 14μl ddH2O and 1μl of worm lysate. The thermocycling conditions of the PCR were 95°C for 2 minutes, followed by 35 cycles of 98°C for 20 seconds, 65°C for 15 seconds for ITS-2 and 70°C for 15 seconds for COX-1, 72°C for 15 seconds and a final extension 72°C for 5 minutes. PCR products were purified with AMPure XP Magnetic Beads (1X) (Beckman coulter, Inc.) using a special magnetic stand and plate according to the protocols described by Beckman coulter, Inc.

A second round PCR was performed using eight forward and twelve reverse barcoded primers (Supplementary Table 2B), avoiding repetitions of the same barcoded primer combinations in different samples. The second round PCR conditions were; 5X KAPA HiFi Fidelity buffer, 10 mM dNTPs, 10 μM barcoded forward (N501 to N508) and reverse (N701 to N712) primers, 0.5 U KAPA HiFi Fidelity Polymerase (KAPA Biosystems,USA), 14 μl ddH2O and 2 μl of first round PCR product as DNA template. The thermocycling conditions of the PCR were 98°C for 45 seconds, followed by 7 cycles of 98°C for 20 seconds, 63°C for 20 seconds, and 72°C for 2 minutes. PCR products were purified with AMPure XP Magnetic Beads (1X) according to the protocols described by Beckman coulter, Inc. A pooled library was prepared from 10 μl of bead purified product from each population and checked with KAPA qPCR library quantification kit (KAPA Biosystems, USA); before being run on an Illumina MiSeq Sequencer using a 500-cycle pair end reagent kit (MiSeq Reagent Kits v2, MS-103-2003) at a concentration of 15 nM with addition 25% Phix Control v3 (Illumina, FC-11-2003).

The MiSeq separates all sequence by sample during post-run processing using recognised indices to generate FASTAQ files. These data were analysed with our own adapted pipeline. Briefly, a NCBI BLASTN search was used to generate and build consensus sequences of the *C. daubneyi* ITS-2 rDNA and mt-COX-1 loci from FASTA files using Geneious Pro 5.4 software (Drummond et al., 2012). Data analysis was performed using Mothur v1.39.5 software (Schloss et al., 2009) and the Illumina Mi-seq standard operating procedures (Kozich et al., 2013). Overall, about 100,000 ITS-2 rDNA and mt-COX-1 reads were generated from the MiSeq data set of 32 *C. daubneyi* populations. In summary, raw paired-end reads were made into contigs, and those that were too long or had ambiguous bases were removed. The sequence data were trimmed to the region amplified by the primers and aligned to the consensus sequence to only include this region. Any sequence data that did not hit with the ITS-2 rDNA and mt-COX-1 consensus sequences were discarded as being trace amplicon contamination.

ITS-2 rDNA and Mt-COX-1 sequence reads were then aligned in Geneious software (Kearse et al., 2012) before removing polymorphisms only occurring once, considered to be artefacts due to sequencing errors (Chaudhry et al., 2015b). The aligned sequences were then imported into the CD-HIT software (Huang et al., 2010) to calculate the frequency of the haplotypes present in each population, and those sequences showing 100% base pair similarity were collapsed into a single haplotype (cd-hit.org/).

### 2.5. Haplotypic diversity of the C. daubneyi mt-COX-1 locus

Within population haplotypic diversity was examined using all of the generated sequences; and between population haplotypic diversity was examined by dividing the number of sequence reads for each haplotype in each population by the number of flukes from which the sequences were derived.

The genetic diversity estimation was calculated from all of the sequences generated using the DnaSP 5.10 software package (Librado and Rozas, 2009). Briefly, sequence polymorphism was estimated through the haplotype frequency (H_f_), variance of haplotype diversity (H_V_), nucleotide diversity (π), the mean number of pairwise differences (k), the number of segregating sites (S) and the mutation parameter based on an infinite site equilibrium model and the number of segregating sites (θ_S_).

### 2.6. Split and network tree analysis of mt-COX-1 haplotypes

A split tree of the mt-COX-1 haplotypes was constructed by HKY+G model of substitution using the Maximum Likelihood method in the SplitTrees4 software (Huson and Bryant, 2006). The program jModeltest 12.2.0 was used to select the appropriate model of nucleotide substitutions for Maximum Likelihood analysis (Posada, 2008). Branch supports were obtained by 1000 bootstraps of the data.

A network tree of 18 mt-COX-1 haplotyes generated this study and other haplotypes at the same locus published on NCBI GenBank was produced based on a neighbour joining algorithm using Network 4.6.1 software (Fluxus Technology Ltd), built on a sparse network with the epsilon parameter is set to zero default. The tree was rooted with the corresponding mt-COX-1 sequence of *Fasciola gigantica* (AJ853848).

### 2.7. Ethical approval

The abattoir based study was approved by the Veterinary Ethical Review Committee of the Royal (Dick) School of Veterinary Studies (VERC 21 14).

## 3. Results

### 3.1. Confirmation of species identity by ITS-2 rDNA sequence

The Illumina MiSeq ITS-2 rDNA data confirmed the species identity of *C. daubneyi* in each of the 32 populations. In all cases, the gross morphology and the size of each worm was typical of *C. daubneyi*, hence the results supported the validity of the rDNA ITS-2 as a genetic marker to identified *C. daubneyi* (Gordon et al., 2013). Two unique haplotypes of the rDNA ITS-2 were identified among the 32 *C. daubneyi* populations, with a single nucleotide polymorphism at position 84 (A/G), when compared with NCBI GeneBank reference sequences derived from Ireland (accession number AB973394.1).

### 3.2. Within population haplotype frequency of mt-COX-locus

A total of eighteen unique haplotypes of the mt-COX-1 locus was identified among the 32 *C. daubneyi* populations, each derived from a single bovine host. In twelve populations, a single haplotype predominated at a frequency of 0.9 – 1.0 (Fig. 1). These comprised of five populations (Pop8, Pop16, Pop19, Pop20, Pop21) of between 10 and 22 parasites that contained a single haplotype, six populations (Pop1, Pop14, Pop25, Pop26, Pop29, Pop31) of between 16 and 35 parasites that contained two haplotypes; and one population (Pop12) of 23 parasites that contained three haplotypes (Table 1, Fig. 1).

**Fig. 1.**
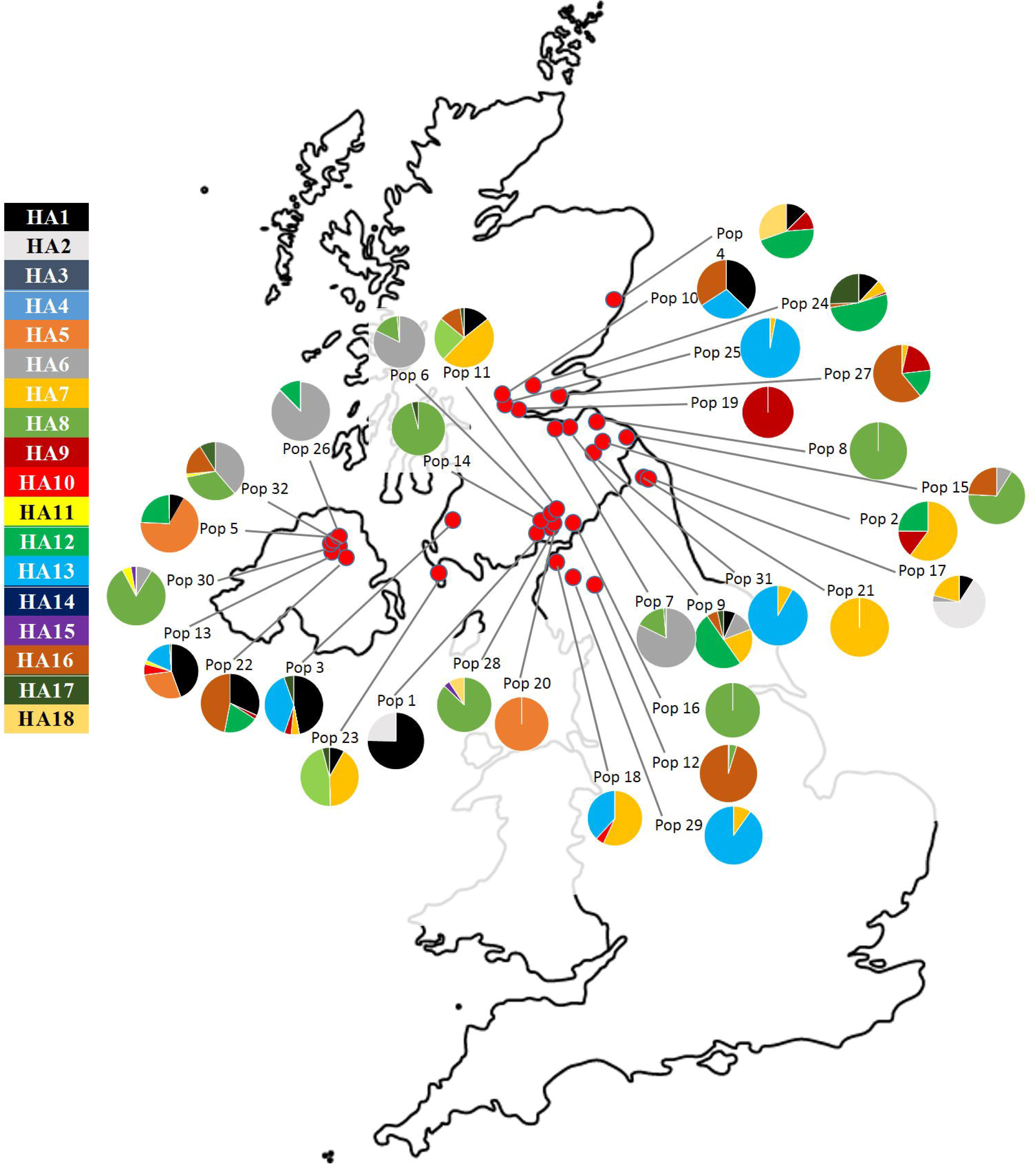
Relative allele frequencies of eighteen individual haplotypes in thirty-two populations. The colours in the pie chart circles indicate the haplotype frequency in each individual population in the map.

In contrast, twenty populations had high frequencies of multiple mt-COX-1 haplotypes (Fig. 1). These comprised of four populations (Pop7, Pop9, Pop13, Pop24) of between 11 and 37 parasites that contained six haplotypes; three populations (Pop3, Pop11, Pop32) of between 31 and 47 parasites that contained five haplotypes; seven populations (Pop4, Pop5, Pop17, Pop22, Pop23, Pop27, Pop30) of between 17 and 43 parasites that contained four 4 haplotypes; and six populations (Pop2, Pop6, Pop10, Pop15, Pop18, Pop28) of between 12 and 28 parasites that contained a maximum of 3 haplotypes (Table 1, Fig. 1).

Overall, the genetic diversity at both haplotype and nucleotide level indicated that the 32 populations were highly diverse, with collapsed overall values of 0.949 and 0.011, respectively (Table 1). There were no patterns in the geographical distribution of the farms from which cattle with more or less diverse populations were consigned to slaughter.

The results would be consistent with a single introduction of *C. daubneyi* infection to some of the farms where the cattle had been grazed during their life history and multiple time introductions to most.

### 3.3. Between population phylogenetic analysis of mt-COX-1 locus

Haplotypes HA8 and HA6 accounted for 35.9% and 13.6% of all of the sequence reads generated per fluke, being present in 10 and 9 populations, respectively. Five haplotypes (HA1, HA7, HA12, HA13 and HA16) accounted for between 5% and 10%; six haplotypes (HA2, HA3, HA5, HA9, HA17, HA18) accounted for between 1% and 5%; and 5 rare haplotypes (HA4, HA10, HA11, HA14, HA15) accounted for less than 1% of the sequence reads generated per fluke. Each of the rare haplotypes was present in between one and four populations.

The split tree of eighteen distinct mt-COX-1 haplotypes in 32 populations (Fig. 2) showed that nine haplotypes (HA1, HA6, HA8, HA9, HA12, HA13, HA15, HA16, HA17) were shared between different *C. daubneyi* populations derived from cattle consigned for slaughter from holdings in SW Scotland/NW England, East Scotland/NE England and Northern Ireland. Three haplotypes (HA2, HA7, HA18) were present in the populations derived from cattle consigned for slaughter from holdings in SW Scotland/NW England and East Scotland/NE England. Three haplotypes (HA5,HA10, HA11) were shared between populations derived from cattle consigned for slaughter from holdings in SW Scotland/NW England and Northern Ireland. The remaining three haplotypes (HA3, HA4, HA14) were only present in a single population derived from cattle consigned for slaughter from a holding in East Scotland, consistent with the emergence, but no spread of these haplotypes.

**Fig. 2.**
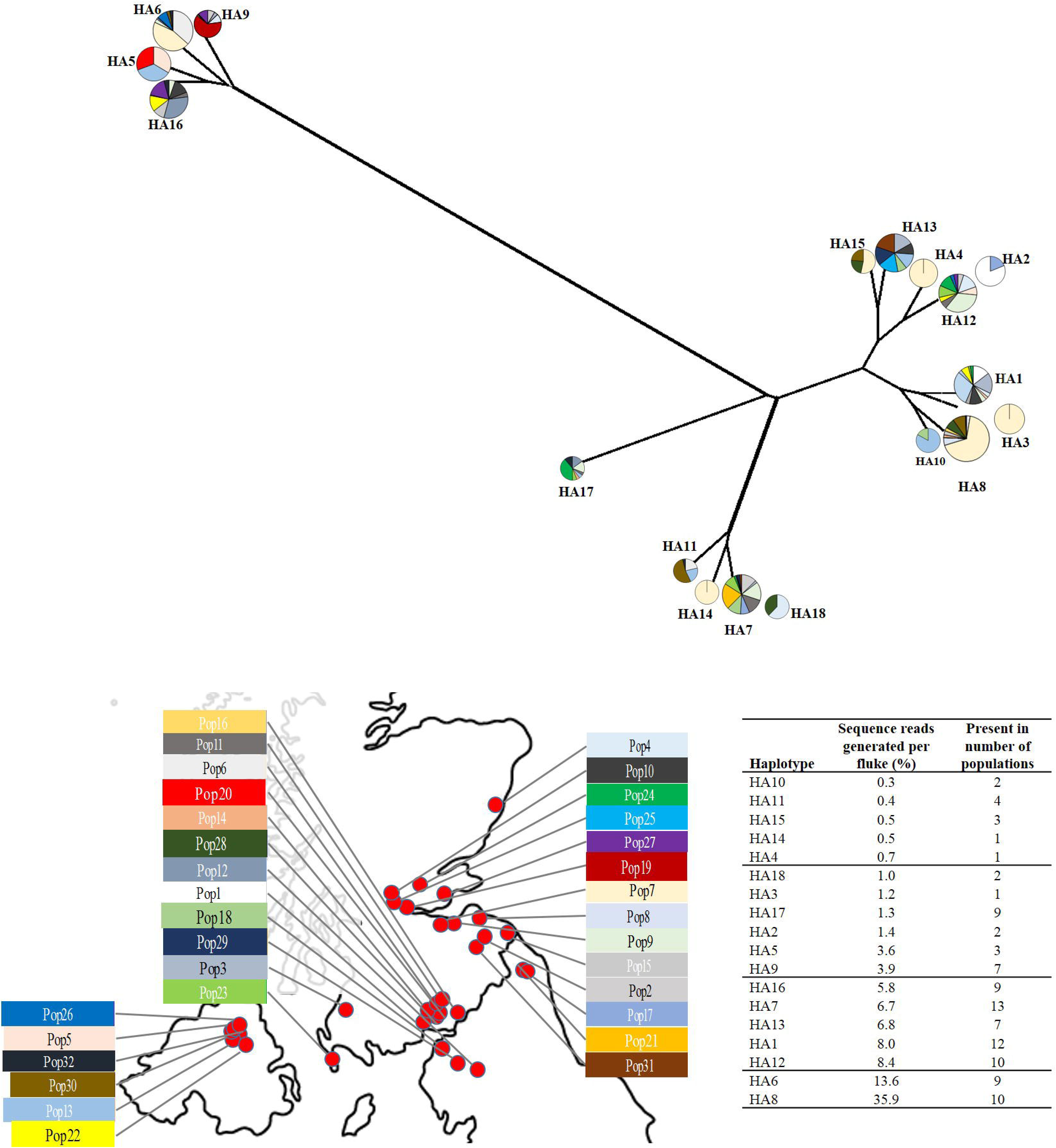
Split tree of the 18 mt-COX-1 haplotypes generated with the median-net method in SplitsTrees4 software (Huson and Bryant, 2006). The colours in the pie chart circles replicate the haplotype frequency in the thirty-two populations indicated on the map. The size of each haplotype circle represents the percentage of sequence reads generated per fluke, shown by rows in the inset table.

The split tree shows all eighteen haplotypes located in three distinct parts of the tree (Fig. 2). Four haplotypes (HA5, HA6, HA9, HA16) are present in a clade I; nine haplotypes (HA1, HA2, HA3, HA4, HA8, HA10, HA12, HA13, HA15) in a clade II; and five haplotypes (HA7, HA11, HA14, HA17, HA18) in a clade III of the network. Fig. 2 shows that the haplotypes are more closely related to each other within, rather than between clades, indicating at least three independent origins of *C. daubney*i infection in the United Kingdom. The split tree showed no clear pattern of geographical distribution of haplotypes, or clades; hence the data are consistent with the spread of haplotypes across several locations after their emergence in the United Kingdom.

Twenty-three unique haplotypes of mitochondrial COX-1 locus were generated in the final filtration for the phylogenetic analyses. The network tree (Fig. 3) shows the 18 *C. daubneyi* mt-COX-1 haplotypes identified in northern United Kingdom, alongside twelve collapsed NCBI GenBank Irish haplotypes, a Spanish haplotype and two collapsed NCBI GenBank haplotypes from South Africa and Zimbabwe. Five of the twelve NCBI GenBank Irish haplotypes were present in SW Scotland/NW England, East Scotland/NE England and Northern Ireland; three in SW Scotland/NW England and East Scotland/NE England; and one in SW Scotland/NW England and Northern Ireland.

**Fig. 3.**
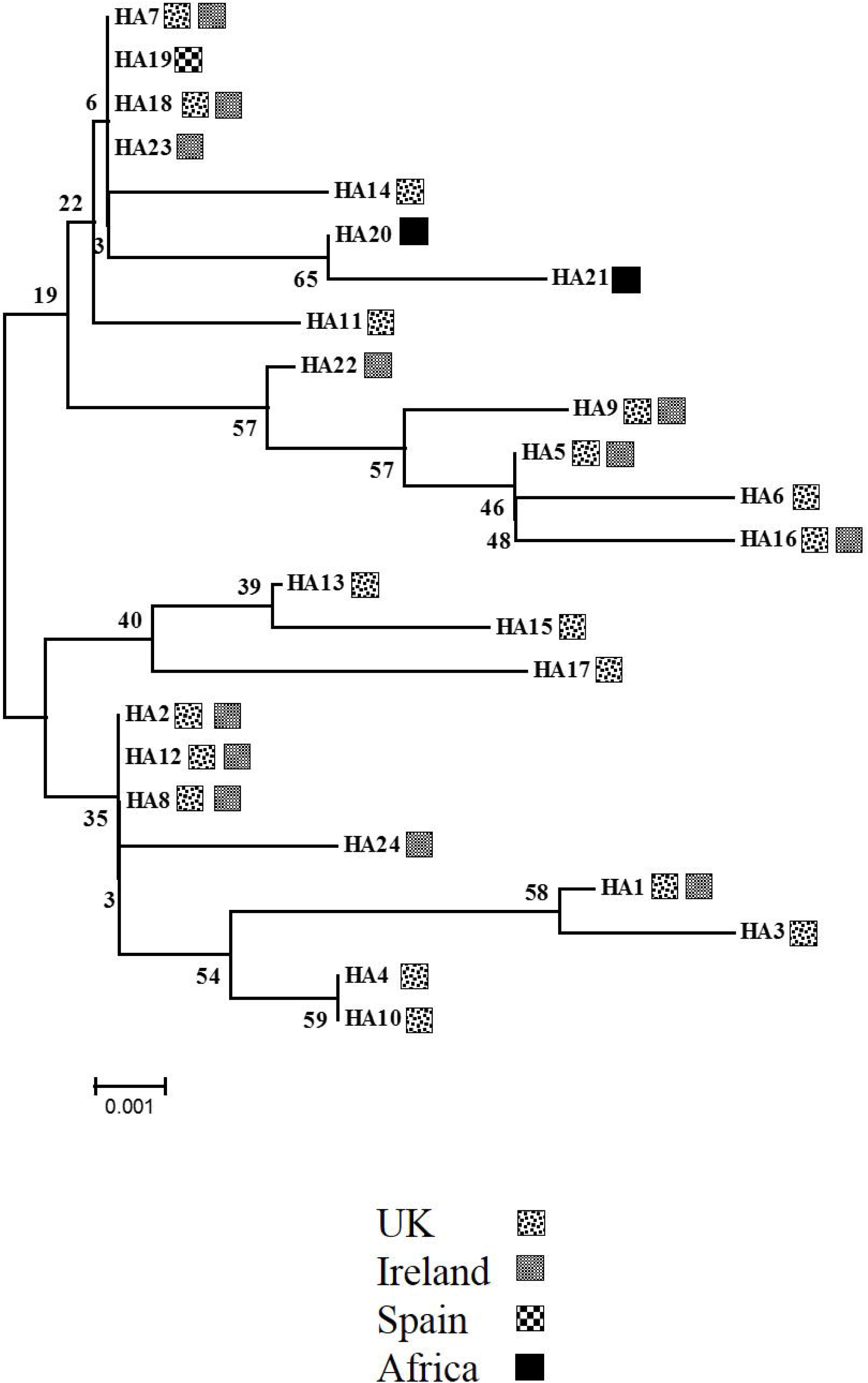
Network tree of the 18 mt-COX-1 haplotypes of the present study aligned with 12 published NCBI GeneBank sequences of *C. daubneyi* from Ireland, Spain and Africa using Geneious Pro 5.4 software (Kearse et al., 2012). Those sequences showing 100% base pair similarity are grouped into a single haplotype using the CD-HIT Suite software (Huang et al., 2010), resulting in 23 unique haplotypes in the final filtration for the phylogenetic analyses. The tree is rooted with the corresponding mt-COX-1 sequence of *Fasciola gigantica*.

## 4. Discussion

There is concern about the probable recent introduction, increased prevalence and potential economic impact of rumen fluke infection of United Kingdom cattle. A study of 339 cattle slaughtered in a Scottish red meat abattoir described a high prevalence of rumen fluke infection in 29% of cattle consigned from Scotland, Northern Ireland and northern England. The study described the geographical distribution of rumen fluke infection, estimated the minimal impact of adult flukes on production, and demonstrated the value of faecal egg counts as a tool to diagnose infection in live animals and to study the epidemiology of the disease (Sargison et al., 2016). Parasites collected from the infected cattle were used as template for the current study.

Deep amplicon sequencing of meta barcoded mt-COX-1 DNA afforded a practical and relatively inexpensive method, when compared to conventional Sanger sequencing (Zintl et al., 2014), to investigate the haplotype diversity between and within populations of *C. daubneyi* rumen flukes derived from individual cattle consigned to slaughter from different locations in SW Scotland/NW England, East Scotland/NE England and Northern Ireland. The Illumina MiSeq platform allows for relatively error-free reads of up to 400 bp (Avramenko, R.W. et al. 2017; Ali et al., 2018), hence the requirement for *de novo* primer design to amplify a 333 bp fragment of the mt-COX-1 locus in this study. The 885 bp fragment (Martínez-Ibeas et al., 2013) used in the Irish cattle phylogenetic study (Zintl et al., 2014) was too large, hence unsuitable for the Illumina MiSeq platform, albeit our primers were designed to amplify a locus within the larger fragment to allow for genomic comparisons. Our method enabled the generation of haplotype sequences from a representative total of 721 parasites, showing the proportions of each haplotype present in populations in individual cattle consigned to slaughter from 32 different farms. The study design allowed the proportions of haplotypes present to be mapped to precise geographical locations from where cattle were consigned to slaughter. However, infection of final hosts is considered to be cumulative, hence our observations may be confounded by both recent infection and burdens established over previous years (González-Warleta et al., 2013), depending on the production system and previous treatments effective against rumen flukes. Some of the cattle in our study may have spent their entire life on a single farm, while others may have had several movements prior to slaughter, hence their *C. daubneyi* burdens at slaughter could have been acquired on other farms, depending on their grazing history and the availability of intermediate snail hosts.

*C. daubneyi* was the only rumen fluke species identified by ITS-2 rDNA sequence data in our study. Data analysis of the full sequence of the ITS-2 rDNA locus only revealed intraspecific variation at a single position, hence this locus was not useful for phylogenetic comparisons. The northern United Kingdom *C. daubneyi* were genetically indistinguishable from Irish populations (Zintl et al., 2014) at the ITS-2 rDNA locus. In contrast, our results showed high levels of haplotype diversity in the mt-COX-1 locus, with 18 unique haplotypes in populations derived from the northern United Kingdom. The nucleotide diversity was also high, highlighting large differences in nucleotide sequences at the same locus. These results are similar to those described in Irish cattle (Zintl et al., 2014), and are consistent with parasite introduction from multiple sources, and a subsequent high mutation rate. This may have important implications for the adaptability of the parasite to potentially favourable factors such as the availability of different intermediate hosts, illustrated by the exploitation of *Galba truncatula* snails by *C. daubneyi* and *Fasciola hepatica* in the United Kingdom (Jones et al., 2015) and changing climatic and management conditions for environmental stages (Skuce et al., 2013); or to potentially adverse factors such as the exposure of parasitic stages to anthelmintic drugs, allowing for the development of anthelmintic resistance.

An aim of the present study was to use novel molecular genetic approaches to gain insight into the emergence and spread of *C. daubneyi* infection in United Kingdom. At farm level, infection may emerge singly or on multiple occasions through animal movement, or translocation of intermediate hosts, or free-living egg, miracidia, cercaria or metacercaria stages. The presence of a single mt-COX-1 haplotype in five populations suggests a single emergence of *C. daubneyi* infection on the farms from which the parasite populations were derived, putatively associated with a combination of geographical isolation and infrequent animal introductions. In contrast, the identification of one population with three unique haplotypes is consistent with multiple introductions, but no subsequent spread of *C. daubneyi* infection, putatively in a cattle-finishing herd with frequent introduction of animals but no onward movements other than to the abattoir. Multiple common mt-COX-1 haplotypes in 26 populations in the absence of geographical clustering of clades is consistent with multiple introductions on the farms from which the parasite populations were derived and subsequent spread of *C. daubneyi* infection. The theoretical alternative explanation of recent expansion of a small divergent established population is unlikely given the high variation in proportions of different haplotypes and the parasite’s reproductive strategy; and due to the absence of reports of rumen flukes in United Kingdom livestock between 1950 (Wilmott, 1950) and 2008 (Foster et al., 2008). Hence, our results illustrate the potential impact of high levels of animal movements in the United Kingdom (Vernon, 2011), for example involving trade in weaned suckler calves and store cattle, on the spread of infection.

In summary, our genetic data provide first insights in the emergence and the spread of *C. daubneyi* infection in United Kingdom. Our findings suggest both single and multiple independent emergence of *C. daubneyi* infection at farm level. Most common mt-COX-1 haplotypes were identified in several populations across a range of geographic locations, highlighting the role of animal movements in the spread of infectious disease. This understanding is relevant to the educational dissemination and implementation of sustainable parasite control strategies. The study provides proof of concept of a method that could be used in the study of host-parasite relationships to examine influences of host movement, animal management, availability of intermediate hosts, and climate change on the epidemiology of parasitic diseases.

## Acknowledgments

We acknowledge Scotbeef Limited who fully funded this project and allowed us to carry out the sampling and collect material. We acknowledge our colleagues who helped with sampling as well as the slaughterhouse workers and the Meat Hygiene Service for making this work possible. Work at the Roslin Institute uses facilities funded by the Biotechnology and Biological Sciences Research Council (BBSRC).

