## Supplementary Table S1 for "A high throughput deep amplicon sequencing method to show the emergence and spread of *Calicophoron daubneyi* in United Kingdom cattle herds"

**Supplementary Table 1:** Number of adult *C. daubneyi* flukes comprising the 32 study populations, each representing the parasite population of a single cattle host consigned to slaughter from a different location. The total number of rumen flukes counted in the forestomach of each host at slaughter is shown.

| **Populations ID** | **Total no of flukes**  **in the forestomach** | **Flukes selected for the present study** | **Date of sample collection** |
| --- | --- | --- | --- |
| Pop1 | 1180 | 11 | 23/09/2014 |
| Pop2 | 103 | 22 | 3/2/2014 |
| Pop3 | 240 | 47 | 27/01/2014 |
| Pop4 | 20 | 20 | 27/01/2014 |
| Pop5 | 2181 | 21 | 27/01/2014 |
| Pop6 | 276 | 27 | 27/01/2014 |
| Pop7 | nr | 37 | 13/01/2014 |
| Pop8 | 20 | 15 | 27/01/2014 |
| Pop9 | nr | 16 | 5/9/2014 |
| Pop10 | 76 | 17 | 13/01/2014 |
| Pop11 | 204 | 32 | 13/01/2014 |
| Pop12 | 66 | 23 | 13/01/2014 |
| Pop13 | 268 | 34 | 13/01/2014 |
| Pop14 | 186 | 35 | 13/01/2014 |
| Pop15 | 44 | 18 | 27/01/2014 |
| Pop16 | 61 | 17 | 6/10/2014 |
| Pop17 | 1169 | 43 | 16/09/2014 |
| Pop18 | 27 | 12 | 16/09/2014 |
| Pop19 | 10 | 10 | 16/09/2014 |
| Pop20 | 249 | 17 | 23/09/2014 |
| Pop21 | 68 | 22 | 16/09/2014 |
| Pop22 | 149 | 11 | 6/10/2014 |
| Pop23 | 25 | 17 | 6/10/2014 |
| Pop24 | 143 | 11 | 6/10/2014 |
| Pop25 | 30 | 24 | 6/10/2014 |
| Pop26 | 736 | 18 | 3/3/2014 |
| Pop27 | 26 | 26 | 3/3/2014 |
| Pop28 | 390 | 28 | 3/2/2014 |
| Pop29 | 64 | 20 | 3/2/2014 |
| Pop30 | 48 | 23 | 25/08/2014 |
| Pop31 | 60 | 16 | 3/2/2014 |
| Pop32 | 1501 | 31 | 25/08/2014 |

nr: not recorded
