## Supplementary Table S2A for "A high throughput deep amplicon sequencing method to show the emergence and spread of *Calicophoron daubneyi* in United Kingdom cattle herds"

**Supplementary Table 2A.** Primer sequences for the amplification of *C. daubneyi* rDNA ITS-2 (AD_For / AD_Rev) and mt-COX-1 (CDX_For / CDX_Rev) forward and reverse primer sets are underlined, N’s are bolded and adapters are in italic format.

| **Sequences (5'-3')** | **Primer name** |
| --- | --- |
| *TCGTCGGCAGCGTCAGATGTGTATAAGAGACAG*GGTGGATCACTCGGCTCGTG | AD_For |
| *TCGTCGGCAGCGTCAGATGTGTATAAGAGACAG***N**GGTGGATCACTCGGCTCGTG | AD_ For-1N |
| *TCGTCGGCAGCGTCAGATGTGTATAAGAGACAG***NN**GGTGGATCACTCGGCTCGTG | AD_ For-2N |
| *TCGTCGGCAGCGTCAGATGTGTATAAGAGACAG***NNN**GGTGGATCACTCGGCTCGTG | AD_ For-3N |
| *GTCTCGTGGGCTCGGAGATGTGTATAAGAGACAG*TTCCTCCGCTTAGTGATATGC | AD_Rev |
| *GTCTCGTGGGCTCGGAGATGTGTATAAGAGACAG***N**TTCCTCCGCTTAGTGATATGC | AD_ Rev-1N |
| *GTCTCGTGGGCTCGGAGATGTGTATAAGAGACAG***NN**TTCCTCCGCTTAGTGATATGC | AD_ Rev-2N |
| *GTCTCGTGGGCTCGGAGATGTGTATAAGAGACAG***NNN**TTCCTCCGCTTAGTGATATGC | AD_ Rev-3N |
| *TCGTCGGCAGCGTCAGATGTGTATAAGAGACAG*ACTTATTTTGTGGGAGTCTTTG | CDX_For |
| *TCGTCGGCAGCGTCAGATGTGTATAAGAGACAG***N**ACTTATTTTGTGGGAGTCTTTG | CDX_For-1N |
| *TCGTCGGCAGCGTCAGATGTGTATAAGAGACAG***NN**ACTTATTTTGTGGGAGTCTTTG | CDX_For-2N |
| *TCGTCGGCAGCGTCAGATGTGTATAAGAGACAG***NNN**ACTTATTTTGTGGGAGTCTTTG | CDX_For-3N |
| *GTCTCGTGGGCTCGGAGATGTGTATAAGAGACAG*CATATTGAATGATAATCGTGACCC | CDX_Rev |
| *GTCTCGTGGGCTCGGAGATGTGTATAAGAGACAG***N** CATATTGAATGATAATCGTGACCC | CDX_Rev-1N |
| *GTCTCGTGGGCTCGGAGATGTGTATAAGAGACAG***NN**CATATTGAATGATAATCGTGACCC | CDX_Rev-2N |
| *GTCTCGTGGGCTCGGAGATGTGTATAAGAGACAG***NNN**CATATTGAATGATAATCGTGACCC | CDX_Rev-3N |
